# Comparison of DNA extraction methods for non-marine molluscs: Is modified CTAB DNA extraction method more efficient than DNA extraction kits?

**DOI:** 10.1101/863167

**Authors:** Sudeshna Chakraborty, Anwesha Saha, N.A. Aravind

## Abstract

Isolation of high molecular weight DNA from gastropod molluscs and its subsequent PCR amplification is considered difficult due to excessive mucopolysaccharides secretion which co-precipitate with DNA and obstruct successful amplification. In an attempt to address this issue, we describe a modified CTAB DNA extraction method that proved to work significantly better with a number of freshwater and terrestrial gastropod taxa. We compared the performance of this method with Qiagen^®^ DNeasy Blood and Tissue Kit. Reproducibility of amplification was verified using a set of taxon-specific primers wherein, modified CTAB extracted DNA could be replicated at least four out of five times but kit extracted DNA could not be replicated. Additionally, sequence quality was significantly better with CTAB extracted DNA. This could be attributed to the removal of polyphenolic compounds by polyvinyl pyrrolidone (PVP) which is the only difference between conventional and modified CTAB DNA extraction methods for animals. The genomic DNA isolated using modified CTAB protocol was of high quality (A260/280 ≥ 1.80) and could be used for downstream reactions even after long term storage (more than two years).

---

Molecular phylogenetics uses a variety of statistical procedures to construct and understand evolutionary relationships among organisms (Sidow and Bowman 1991). These relationships are crucial in addressing questions related to delimitation of species, divergence time estimation, and study of fossils among other applications of phylogenetics (Thorne and Kishino 2002; Giribet 2003; Yang and Rannala 2010). There was an unprecedented growth in this field with the advent of various sophisticated techniques namely cloning, enzymatic amplification (Lio and Goldman 1998), whole genome sequencing (Dowell 2008), etc. Despite being diverse in nature, a common and the primary step in each of these methods is extraction of high molecular weight DNA. It is also arguably the most pivotal step since the accuracy and quality of the final results depend to a large extent on the quality of the DNA.

Several genomic DNA extraction methods have been described for animals which include both the manual methods (Cheung *et al*. 1993; Winnepenninckx *et al*. 1993; Aljanabi and Martinez 1997; Yue and Orban 2005) and commercially available extraction kits. However, in the case of gastropod molluscs and bivalves, the presence of excessive slime (mucopolysaccharides) is detrimental due to its amplification inhibiting capacity. It co-precipitates with DNA, inhibiting the enzyme activity during PCR (Winnepenninckx *et al*. 1993; Sokolov 2000; Popa *et al*. 2007). While much has been written about bivalves and other marine molluscs in this regard (Aranishi and Okimoto 2006; Popa *et al*. 2007; Pereira *et al*. 2011), their terrestrial and freshwater counterparts have received very little attention. In this communication, we provide a detailed description of a genomic DNA extraction method that proved to work well with many terrestrial as well as freshwater gastropod families producing high-quality DNA. The extracted DNA from both fresh and old samples gave consistent results when used for PCR. We have also compared its performance with Qiagen DNeasy^®^ Blood and Tissue Kit since it is one of the most widely used commercially available kits for DNA extraction.

This study included 23 representative individuals belonging to 13 families, 18 genera and 23 species collected from the Western Ghats of India (Table S2). All specimens were thoroughly washed with absolute ethanol initially and kept immersed in it. For the next one week, ethanol was changed once every two days since it became turbid due to slime discharge by the animal. Additionally, ethanol was changed every time it turned turbid or yellowish over time due to slime discharge to ensure proper preservation of the animal. Prior to DNA extraction, about 20mg of tissue (whole body excluding the shell if the animal was tiny) was kept immersed in twice the volume of absolute ethanol for another two days to remove any remnant slime present on the tissue.

## Modified CTAB method

The alcohol-soaked tissue was crushed slightly using a pestle and transferred to 400 µL TE buffer for softening before removing excess alcohol using Kimtech® Kimwipes and incubated at room temperature (25 – 30°C) with mild shaking at 400 rpm for an hour. In the case of old samples (a few years old), if not preserved properly, the slime hardens and resembles the tissue. This makes it difficult to differentiate between the two. Therefore, the treatment with TE buffer becomes crucial since it dissolves the slime entirely leaving behind only the tissue. After incubation, the softened tissue was immersed in 400 µL of CTAB buffer pre-heated to 60°C. The tissue was then subjected to mechanical disruption using a bead beater post adding 20 µL Qiagen® Proteinase K and a silica bead, and kept for overnight digestion at 60°C. The suspension was extracted with 400 µL of chloroform: isoamyl alcohol (24:1) thrice at 12,000 rpm for 5 min at 4°C. The supernatant was carefully separated to ensure the white layer remained undisturbed and precipitated with 800 µL of absolute ethanol in the presence of 40 µL 3M Sodium acetate at 12,000 rpm for 10 minutes at 4°C. The pellet was washed again with 200 µL 70% ethanol, air dried, and re-suspended in TE Buffer for long-term storage. The components and corresponding quantities of all the reagents used are summarized in Table 1. All reagents were autoclaved after preparation except lysis buffer in which CTAB was added after autoclaving. Since RNA was not found to hinder with the amplification process unlike polyphenols, no RNase treatment was done.

**Table 1:** List of reagents used in modified CTAB method.

| S.No. | Reagent | Components | Quantity |
| --- | --- | --- | --- |
| 1 | 1M Tris-HCl | Tris Base | 121.1g |
|  |  | HCl | Make pH 8.0 |
|  |  | Distilled Water | Adjust volume to 1L |
| 2 | 0.5M EDTA disodium dihydrate | EDTA disodium dihydrate | 186.1g |
|  |  | Sodium Hydroxide | Make pH 8.0 |
|  |  | Distilled Water | Adjust volume to 1L |
| 3 | 5M Sodium chloride | Sodium Chloride | 292.2g |
|  |  | Distilled Water | Adjust volume to 1L |
|  |  | Tris-HCl (pH 8.0) | 100mM |
| 4 | Lysis Buffer | Sodium Chloride | 1.4M |
|  |  | EDTA disodium dihydrate | 20mM |
|  |  | Hexadecyltrimethylammonium bromide (CTAB) | 2% w/v |
|  |  | Polyvinyl pyrrolidone | 2% w/v |
| | | β-mercaptoethanol | 0.2% v/v |
| 5 | 3M Sodium Acetate | Sodium acetate trihydrate | 102.025g |
|  |  | Glacial acetic acid | Adjust pH to 5.2 |
|  |  | Distilled Water | Adjust volume to 250mL |
| 6 | TE Buffer | 1M Tris Cl | 0.5mL |
| | | 0.5M EDTA | 100μL |
|  |  | Distilled Water | 50mL |

## Qiagen DNeasy® Blood and Tissue Kit (Cat No. 69504)

Extraction of DNA was performed according to the manufacturer’s protocol with the following minor modifications. To begin with, samples were kept for overnight digestion since foot tissues of snails do not get entirely lysed within a few hours. Additionally, in the final step, DNA was eluted twice by adding 50µL of Buffer AE instead of 200µL to obtain a higher final concentration of DNA.

The extracted DNA samples were measured quantitatively using a spectrophotometer. All the successfully extracted samples (A260/280 ≥ 1.80) were amplified with Qiagen^®^ Taq PCR core kit (Cat No. 201225) using 28S and 16S rRNA primers (Table S1) and sequenced. Both modified CTAB and kit extracted DNA, though comparable in quality (Table S3), differed significantly in their consistency of repeated amplification. Despite showing a high A260/280 value, the results of enzymatic amplification of kit extracted DNA could not be reliably reproduced. For instance, if a kit extracted DNA sample was amplified five times using a set of taxon-specific primers, consistent amplification was observed not more than twice. On the other hand, if the same sample was extracted using CTAB buffer, it amplified at least four out of five times. Additionally, most samples extracted using DNA extraction kit either could not be amplified or gave very poor quality sequences. This efficiency of the CTAB extraction protocol was consistent for both freshly extracted (few days old) as well as stored DNA samples (over 5 years old). A possible reason for this hindrance could be the presence of polyphenols.

Algal association has not only been studied for marine gastropods (Geiselman and McConnell 1981; Steinberg 1988) but the herbivorous and algivorous nature of both terrestrial and freshwater snails are also well documented (Mason 1970; Butler 1976; CARTER *et al*. 1979; Brönmark 1989; Speiser and Rowell-Rahier 1991; Vaughn *et al*. 1993; Martin 2000; March *et al*. 2002; Chase and Knight 2006; Fink and Von Elert 2006). We have observed diatoms in the gut of *Cremnoconchus* sp. (Littorinidae). The preference of land snails to thrive on extremely moist areas rich in algae and considerable algal growth on the shells of most freshwater snails further strengthen this argument. This makes the presence of polyphenols in the snail tissues all the more likely. In the case of bigger animals, when only foot tissue was used for DNA extraction, the consistency of amplification was comparatively better than smaller animals where the entire body had to be used (pers. obs.). Although the quality of extracted DNA was similar in both the cases, successful amplification and reproducibility of obtained results was the hurdle. The quality of DNA (A260/280) was not always indicative of its amplification capabilities. This is evident from chromatograms in fig. 1 and 2 which clearly depict the quality difference between sequences of CTAB and kit extracted DNA respectively. Additionally, a comparison of average quality of sequences obtained by amplification of CTAB and kit extracted DNA is illustrated in fig. 3. This could be attributed to the presence of increased amounts of polyphenols in the gut compared to the external foot tissue which further validates this hypothesis.

**Fig 1.**
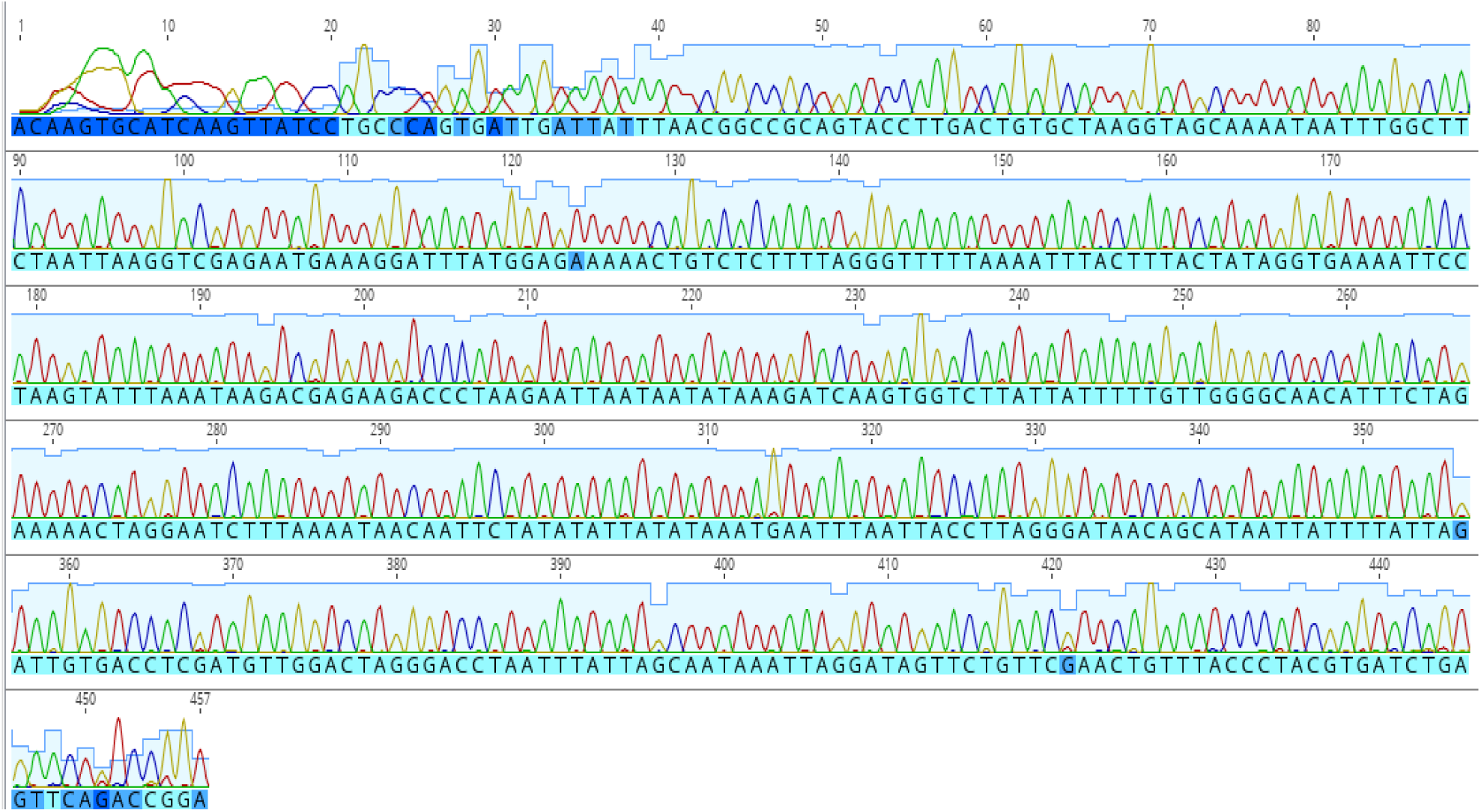
Chromatogram depicting 16S gene sequence quality in modified CTAB extracted DNA.

**Fig 2.**
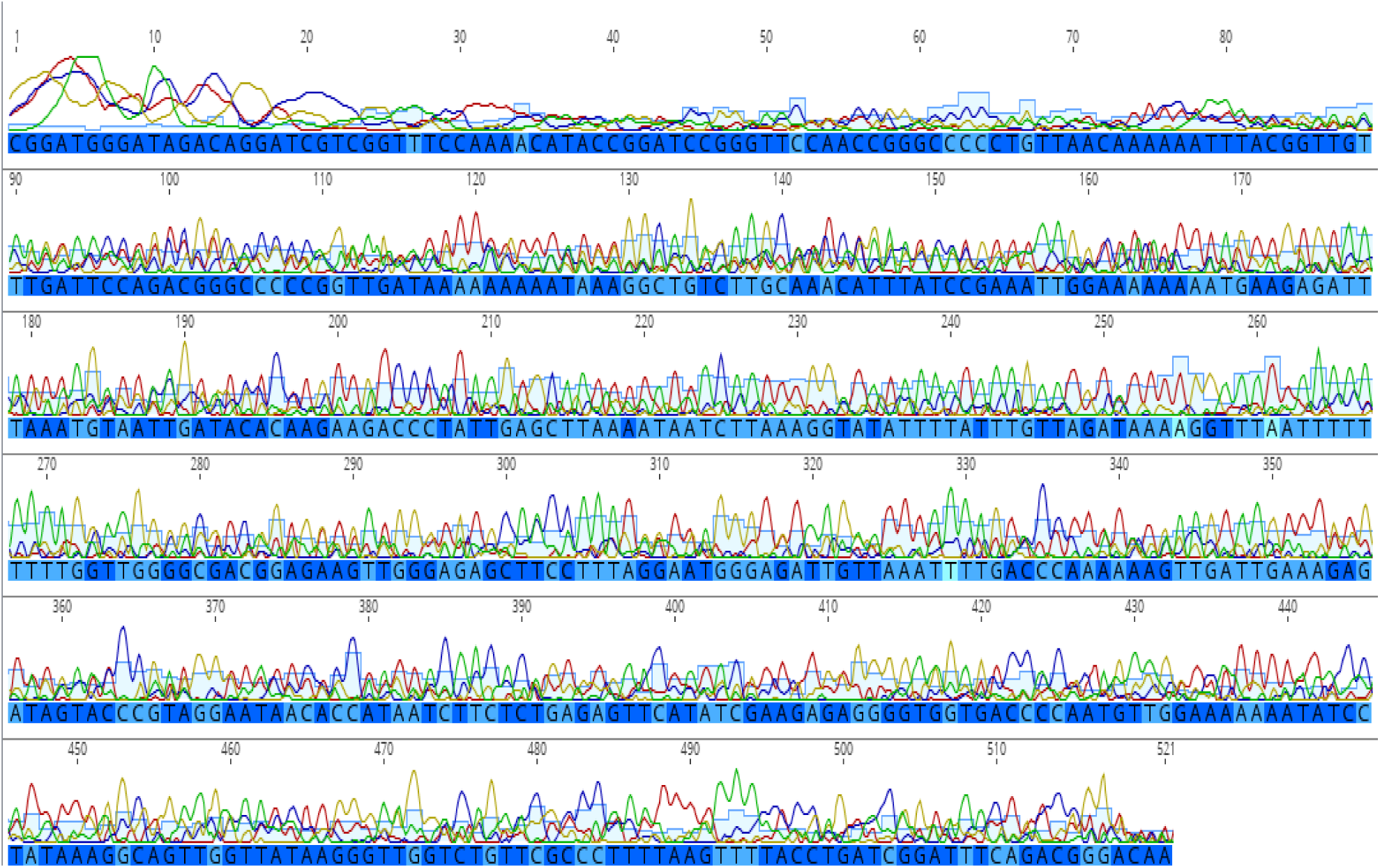
Chromatogram depicting 16S gene sequence quality in Qiagen^®^ DNeasy Blood and Tissue Kit extracted DNA.

**Fig 3.**
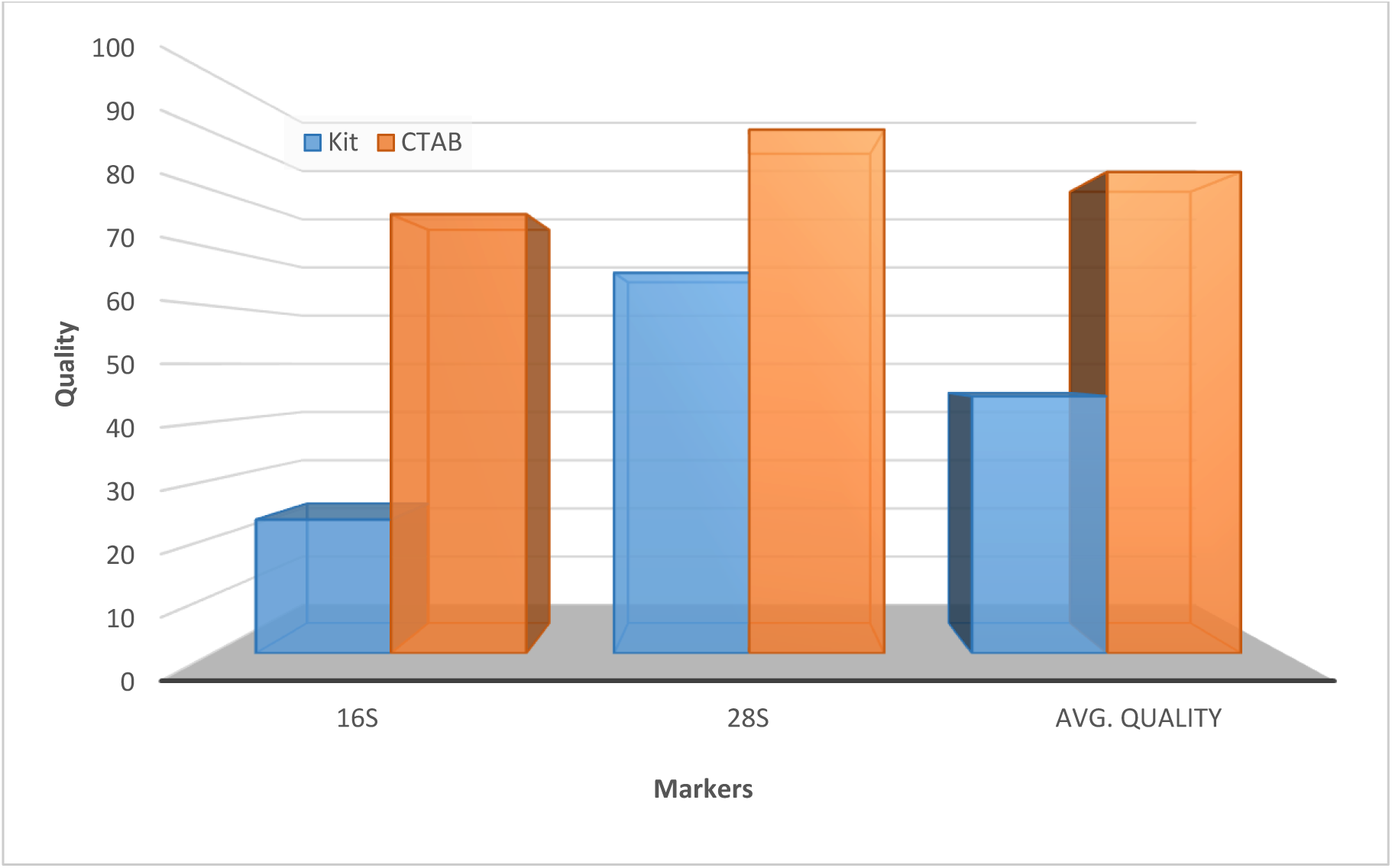
Comparison of average sequence quality between CTAB and Qiagen® DNeasy Blood and Tissue Kit extracted amplified genomic DNA

Most studies that specifically deal with DNA extraction from molluscs, almost always mention the problems of co-precipitation of mucopolysaccharides with DNA (Sokolov 2000; Skujiene and Soroka 2003; Huelsken *et al*. 2011; Poonam *et al*. 2013; Jaksch *et al*. 2016) but exclude the polyphenolic impurities. The prominent association of algal cells and consecutively polyphenols with snails has, invariably, been overlooked. We attribute the hindrance of inconsistent amplification to the presence of polyphenols since the main difference between the modified CTAB extraction protocol used and the conventional method employed for animal DNA extraction was the inclusion of polyvinylpyrrolidone (PVP). Utility of PVP in plant DNA extraction has been extensively studied and its role has been identified to be the removal of polyphenols (John 1992; Porebski *et al*. 1997). Efficacy of PVP utilization in genomic DNA extraction of marine gastropods has also been reported earlier (Williams *et al*. 2003). The same proved to be true across several non-marine gastropod families included in the study. The differences between the methods discussed here have been summarized in Table 2.

**Table 2:**
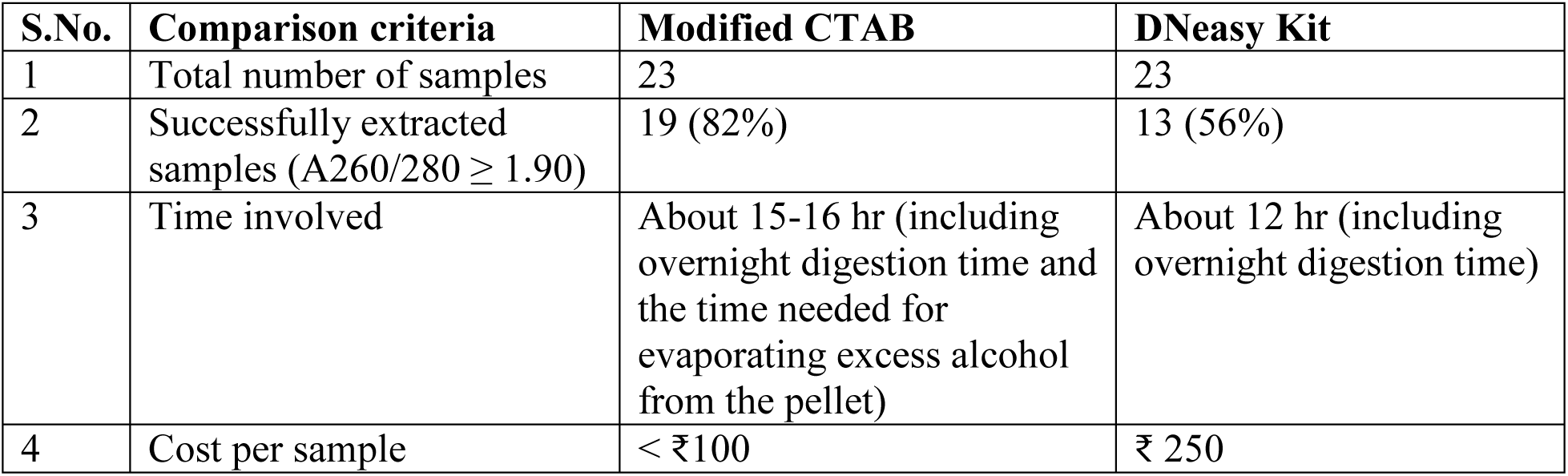
Comparison between the two methods of DNA extraction.

| S.No. | Comparison criteria | Modified CTAB | DNeasy Kit |
| --- | --- | --- | --- |
| 1 | Total number of samples | 23 | 23 |
| 2 | Successfully extracted samples ($A_{260}/A_{280} \geq 1.90$) | 19 (82%) | 13 (56%) |
| 3 | Time involved | About 15-16 hr (including overnight digestion time and the time needed for evaporating excess alcohol from the pellet) | About 12 hr (including overnight digestion time) |
| 4 | Cost per sample | < ₹100 | ₹ 250 |

The conventional CTAB DNA extraction method was also implemented for extraction of genomic DNA from gastropod molluscs wherein the lysis buffer used consisted of CTAB, and β-mercaptoethanol (data not included). It was treated with sodium chloride (NaCl) post overnight digestion with the lysis buffer. But the quality and quantity of DNA obtained was extremely poor. As an additional measure, the extracted DNA was kept in isopropanol overnight to improve quality. Amplifiable genomic DNA still could not be extracted. This scenario changed only with the inclusion of PVP in the lysis buffer. On addition of PVP, the second overnight treatment with isopropanol was not required. The only difference between the two protocols employed, reinforce the crucial part played by PVP in removal of polyphenolic impurities rendering the extracted DNA suitable for downstream reactions.

An added advantage of this method is the requirement of only a small amount of foot tissue and not an internal organ like liver in case of bigger animals. Unless there is a requirement for extensive morphological and anatomical studies, our modified protocol opens options for non-lethal sampling wherein only a small part of the foot tissue is cut without the necessity of whole animal preservation. This method of sampling is especially handy when working in protected areas since disturbance is minimal.

A few studies have mentioned a possible partial degradation of plant DNA in the presence of CTAB during extraction (Fang *et al*. 1992; Rowland 1993). However, in the case of snails, no such issue was encountered. On the contrary, CTAB extracted DNA was found to be very stable and could be used for amplification even after about two years of its extraction. Another possible shortcoming of the CTAB method could be the time and labor involved, given specialized commercially available kits like E.Z.N.A^®^ Mollusc DNA kit which require lesser time and energy. However, the aforementioned kit is considerably expensive and overlooks the fact that polyphenolic compounds also hinder amplification and not mucopolysaccharides alone. Apart from being a relatively cheap option, our modified CTAB protocol saves the cost and effort of repeated amplification which becomes extremely important in the long run. The difference in the cost of modified CTAB extraction and kit extraction of DNA in conjunction with its added efficiency may be useful for labs with access to limited resources. The chemicals used in the CTAB extraction method are readily available as opposed to the highly specialized kits which are not always an easy option especially for projects with limited funding. In summary, this protocol can be used across non-marine molluscan taxa to successfully obtain high-quality genomic DNA without hindrance by polysaccharide or polyphenolic impurities rendering a high degree of success and consistency to enzymatic amplification reactions.

## Acknowledgements

Authors would like to thank Department of Science and Technology, Government of India (Grant No. EMR_2015/000199/v1/21385) for funding. We would also like to thank Dr. Suzanne Williams, Dr. Praveen Karanth, Dr. Maitreya Sil for help in developing the protocol.

## Supplementary information

**Table S1:**
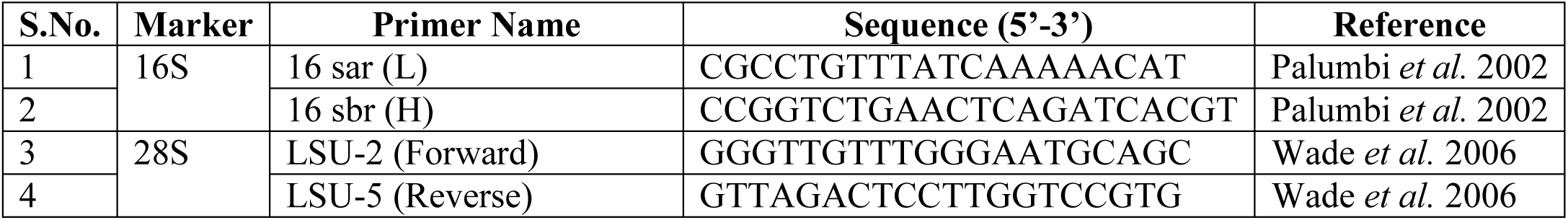
List of primers used.

**Table S2:** List of samples used in the study and the quality (A260/280 value) of extracted DNA.

| S.No. | Habitat | Group | Family | Genus | Quality of DNA (A260/280) |  |
| --- | --- | --- | --- | --- | --- | --- |
|  |  |  |  |  | CTAB | Kit |
| 1 | Freshwater | Caenogastropoda | Viviparidae | *Bellamya bengalensis* | 1.95 | 1.92 |
| 2 | Freshwater | Caenogastropoda | Viviparidae | *Bellamya* sp. | 2.12 | 2.10 |
| 3 | Freshwater | Caenogastropoda | Planorbidae | *Planorbis* sp. | 1.88 | 1.94 |
| 4 | Freshwater | Caenogastropoda | Planorbidae | *Indoplanorbis exustus* | 2.14 | 2.12 |
| 6 | Freshwater | Caenogastropoda | Pachychillidae | *Paracrostoma* sp. | 2.05 | 1.86 |
| 7 | Freshwater | Caenogastropoda | Thiaridae | *Thiara* sp. | 2.00 | 1.89 |
| 8 | Freshwater | Caenogastropoda | Pleuroceridae | *Paludomas* sp. | 2.03 | 2.02 |
| 9 | Terrestrial | Caenogastropoda | Cyclophoridae | *Cyclophorus* sp. | 2.01 | 1.96 |
| 10 | Terrestrial | Caenogastropoda | Cyclophoridae | *Cyclophorus* sp. | 2.11 | 1.94 |
| 11 | Terrestrial | Caenogastropoda | Cyclophoridae | *Theobaldius* sp. | 2.03 | 1.85 |
| 12 | Terrestrial | Caenogastropoda | Pupinidae | *Tortulosa* sp. | 1.53 | 1.36 |
| 13 | Terrestrial | Pulmonata | Camaenidae | *Trachia crassicostata* | 1.81 | 1.62 |
| 14 | Terrestrial | Pulmonata | Camaenidae | *Beddomea calcadensis* | 1.96 | 1.87 |
| 15 | Terrestrial | Pulmonata | Camaenidae | *Trachia vittata* | 1.92 | 1.77 |
| 16 | Terrestrial | Pulmonata | Subulinidae | *Glessula* sp. | 1.99 | 2.03 |
| 17 | Terrestrial | Pulmonata | Subulinidae | *Glessula* sp. | 2.02 | 1.42 |
| 18 | Terrestrial | Pulmonata | Helicarionidae | *Eurychlamys* sp. | 2.01 | 1.96 |
| 19 | Terrestrial | Pulmonata | Helicarionidae | *Eurychlamys* sp. | 2.05 | 1.99 |
| 20 | Terrestrial | Pulmonata | Cerastidae | *Rachis* sp. | 1.99 | 2.04 |
| 21 | Terrestrial | Pulmonata | Cerastidae | *Rachis* sp. | 2.02 | 2.01 |
| 22 | Terrestrial | Pulmonata | Camenidae | *Chloritis* sp. | 2.05 | 2.00 |
| 23 | Terrestrial | Pulmonata | Corillidae | *Corilla* sp. | 1.92 | 1.68 |

